# XPF mediates 3’ flap processing for FEN1-independent Okazaki fragment maturation

**DOI:** 10.1101/2025.10.24.684453

**Authors:** Kejiao Li, Feng Yang, Yingying Wang, Guojun Shi, Yao Yan, Yi Lei, Yixing Wang, Main Zhou, Haitao Sun, Li Zheng, Binghui Shen

**Author notes:** Correspondence should be addressed to: Haitao Sun, Li Zheng, Binghui Shen.

## Abstract

Okazaki fragment maturation (OFM), the process that removes RNA-DNA primers, is a major source of DNA replication stress and mutations. It involves Polδ-mediated DNA strand displacement synthesis that produces 5’ flaps, FEN1-mediated 5’ flap cleavage, and LIG1-catalyzed nick ligation. Recently, we discovered that under stress conditions, yeast cells convert 5’ flaps into 3’ flaps, which are degraded by 3’ flap nucleases to produce ligatable DNA nicks. However, little is known about this stress-induced, 3’ flap-based OFM in human cells. Here, we report that 3’ flaps frequently form in various human cancer cells, and that FEN1 deficiency significantly enhances 3’ flap levels. XPF1 is recruited to the replication forks in FEN1 mutant or FEN1-chemically inhibited cells. Notably, XPF deficiency or inhibition in those defective cells leads to accumulation of 3’ flaps, replication-related DNA strand breaks, and unique mutation signatures. Furthermore, XPF and FEN1 inhibitors show synergistic effects in killing human cancer cells. In summary, we demonstrate that 3’ flap-based OFM is an important alternative of 5’ flap-based OFM in mammalian cells. XPF is a key nuclease to degrade 3’ flaps and complete OFM for survival. Targeting this compensatory mechanism could provide new therapeutic strategies to selectively impair cancer cell survival under pre-existed replication stress.

**Highlights:**

- XPF is recruited to DNA replication forks in mouse or human cells of functional deficiency of FEN1 due to genetic mutations or chemical inhibition.
- XPF gene deficiency or chemical inhibition results in accumulation of 3’ flaps and replicative DNA single-stranded breaks.
- Functional deficiency of XPF and FEN1 displays a synergy in inducing genome instability and cell death in human cancer cells.

## Introduction

Replication stress is a major source of genome stability and the driving force behind the initiation and malignant progression of tumor cells (1–3). On the other hand, replication stress, a hallmark of cancer cells, has been considered as the Achilles’ heel of cancer therapy such as radio-and chemotherapy because excessive replication stress induce cellular senescence or cell death (4–6). During different DNA metabolic processes including DNA replication and repair, various intermediate DNA structures are generated, and their timely and proper resolution is critical for maintaining genomic integrity (7). Notably, during lagging strand DNA synthesis, which occurs discontinuously, millions of Okazaki fragments are produced per cell cycle in human cells (8,9). Thus, the process of transforming these fragments into a continuous lagging DNA strand, known as Okazaki fragment maturation (OFM), is essential for genome integrity and cell survival (8,10,11). Defects in OFM lead to the accumulation of DNA breaks, resulting in replication stresses that may cause genome instability or cell death (12–14).

Dynamic formation and nucleolytic degradation of single-stranded flap structures are key processes in OFM (11,15,16). During lagging strand DNA synthesis under normal physiological conditions, DNA polymerase δ (Pol δ) extends the nascent DNA from the Pol α-synthesized RNA-DNA primer. Continuous DNA synthesis then displaces the RNA-DNA primer of the downstream Okazaki fragment, producing a 5’ single-stranded flap structure. At this stage, Pol δ performs strand displacement synthesis, generating a single-strand flap structure (17). This RNA-DNA 5’ flap is subsequently recognized and cleaved by flap endonuclease 1 (FEN1), and the resulting nick is sealed by DNA ligase I (LIG1) to generate a continuous and fully ligated DNA strand (18,19). If a Pol α-incorporated error is present downstream of the 5’ flap, MutSα recognizes the mismatch and stimulates FEN1 to use its exonuclease activity to remove the error (20). Therefore, the 5’ flap based OFM is considered a highly faithful process. In this 5’ flap-based OFM, FEN1 plays a central role. FEN1 mutations impairing its flap endonuclease or exonuclease activity or abolishing its dynamic interactions, lead to defects in 5’ flap cleavage and accumulation of unligated Okazaki fragments (12–14). The single strand breaks (SSBs) may be further converted into DNA double strand breaks (DSBs), which are the most mutagenic and lethal type of DNA damage. In yeast, deletion of RDA27 (the yeast homolog of FEN1) causes cell death at the restrictive growth temperature of 37°C (12–14), and FEN1 deficiency or mutations in mammalian cells also result in cell growth defects and cell death (21,22). These findings demonstrate the essential role of FEN1-mediated 5’ flap cleavage in OFM.

Intriguingly, a subset of *rad27* knockout yeast cells can escape the lethal effects of restrictive temperature (11). Subsequent studies revealed that this occurs because *rad27* knockout cells grown at 37°C activate DUN1 signaling, which transforms 5’ flaps into 3’ flaps (11). The 3’ flap nuclease activity of Pol δ cleaves the 3’ flap to generate ligatable DNA nick for OF ligation. However, in certain 3’ flaps, the 3’ end may fold back or anneal to nearby ssDNA regions and extend itself, forming secondary structures that are resistant to Polδ-mediated degradation. It is also possible that other 3’ flap nucleases can recognize and cleave these 3’ flaps. However, if such 3’ flaps escape degradation, they can result in DNA duplications containing an internal sequence between two duplication units. Inhibition of DUN1 blocks 3’ flap formation and the generation of internal tandem duplications (ITDs), such as *pol3*-ITD. These observations in yeast indicate that DUN1-induced 3’ flap OFM serves as an important alternative to the canonical 5’ flap-based OFM during DNA replication under stress conditions. However, little is known about how 5’ flaps are converted into 3’ flaps, or which 3’ nucleases, in addition to the 3’ flap nuclease activity of Polδ, catalyze 3’ flap removal during OFM. Furthermore, ITDs like those observed in *rad27* knockout yeast cells frequently occur in human cancers (23–26). Suggesting that 3’ flap-based OFM is conserved in human cells and may contribute to cancer initiation and progression by inducing mutations and promoting cell survival. Nevertheless, the extent by which 3’ flap-based OFM contributes to DNA replication in human cells remains unclear.

In the current study, we screened for nucleases that genetically interact with RAD27 (showing synthetic growth defects or lethality) in yeast. We identified RAD1, the yeast homolog of XPF (encoding ERCC4) (27), as a candidate enzyme responsible for 3’ flaps cleavage. Previous studies have shown that, as a member of the XPF/MUS81 family, XPF forms a complex with ERCC1 and consists of one catalytic and one noncatalytic subunit, exhibiting endonuclease activity toward various 3’ flap and fork DNA structures (27,28). We demonstrate that XPF is not associated with replication forks in wild-type cells, but it is recruited to replication fork when FEN1 is mutated or inhibited. XPF knockout or inhibition results in the accumulation of 3’ flaps in human cancer cells, particularly under FEN1 deficient of inhibited conditions. Furthermore, XPF knockout or inhibition displays synergistic effects with FEN1 inhibitor in killing cancer cells. Our study reveals a critical molecular mechanism by which cancer cells overcome replication stress through an error-prone 3’ flap-based OFM pathway. Targeting this compensatory mechanism may provide novel therapeutic strategies to selectively impair cancer cell survival under replication stress.

## Results

### Identification of helicases and 3’ flap nucleases potentially involved in 3’ flap OFM

To define 3’ flap-based OFM in yeast and human cells, we sought to answer two fundamental questions: how unprocessed 5’ flaps are converted into 3’ flaps and which 3’ flap nucleases catalyze the nucleolytic degradation of 3’ flaps. We previously demonstrated that activation of Dun1, the homolog of human CHK2 (29), is vital for the formation of 3’ flaps, the production of alternative duplications, and the generation of revertants in *rad27Δ* yeast cells (11). It is plausible that the helicases and nucleases required for 3’ flap formation and processing are physically associated with the Dun1 network and are phosphorylated by this signaling cascade. We therefore surveyed the Saccharomyces genome database (30) for helicases and nucleases that physically interact with Dun1 and its activators Mec1 and Rad53. These include the helicases Pif1, Sgs1, and Srs2 and the nucleases Rad1, Mus81, Mre11, and Sae2 (Supplementary Figure S1).

If a helicase or a nuclease is important for 3’ flap based OFM that backs up Rad27-centered 5’ flap based OFM, it should genetically interact with RAD27. Thus, mutant yeast cells are expected to show either synthetic growth defects (SGD) or synthetic lethality (SL) due to the combined deficiency of both 5’ flap-and 3’ flap-based OFM pathways. Supporting this hypothesis, our survey of the yeast genetics database (30) indicated that *rad27Δ* yeast cells exhibited synthetic growth defects (SGD) with Dun1-associated helicase *pif1Δ*, and *rad27Δ* displayed synthetic lethality (SL) with the strain deleted of either of Dun1-associated helicases and nucleases including *sgs1Δ, srs2Δ, rad1Δ, mus81Δ, mre11Δ, or sae2Δ* (31). We previously showed that the *rad27Δ* growth defect phenotype could be rescued by the *pol3-ITD* mutation, which limits 5’ flap formation (11). Therefore, if the observed SGD or SL is driven by the 5’ flap accumulation during OFM, the *pol3-ITD* mutation should be able to rescue these phenotypes as well. To verify this, and to determine which SGD or SL phenotype can be rescued by *pol3-ITD*, we crossed the *rad27Δ* or *rad27Δ*-*pol3-ITD* strains with yeast strains carrying deletions in *sgs1Δ, srs2Δ, rad1Δ, mus81Δ, mre11Δ*, or *sae2Δ*. We performed spore assay or random spore analyses to evaluate the survival of yeast strains with different genotypes and confirmed the SGD or SL phenotypes (Supplementary Figure2, Table S1-S6). The *pol3-ITD* mutation partially rescued the SGD of *rad27Δ* with *pif1Δ* (Supplementary Figure S2A) and fully rescued the SL of rad27Δ with sgs1Δ, rad1Δ, or mus81Δ, but not with mre11Δ (Table 1, Supplementary Table S1-S6). To further confirm that pol3-ITD indeed suppresses the SL phenotype, we conducted yeast spot assays, which demonstrated that *pol3-ITD*::*rad27Δ*::*rad1Δ* or *pol3-ITD*::*rad27Δ*::*mus81Δ* yeast strain had similar normal growth rate as the WT strain (Supplementary Figure S2B). Taking together, our yeast genetic analysis suggests that Pif1, Sgs1, Rad1, and Mus81 function as back up factors for Rad27-mediated OFM, and they may also participate in 3’ flap-based OFM in mammalian cells.

**Table 1:** Synthetic growth defect or lethality between rad27Δ and 3’ helicases and 3’ nucleases.

| Category | Yeast Gene | Human Homologues | wt Pol3 | Pol3-ITD |
| --- | --- | --- | --- | --- |
| Helicase | SGS1 | RECQs | SL* | normal |
|  | SRS2 | FBH1 | SL | SL |
|  | PIF1 | PIF1 | SGD** | normal |
| 3' Nuclease | RAD1 | XPF | SL | normal |
|  | MUS81 | MUS81 | SL | normal |
|  | MRE11 | MRE11 | SL | SL |
|  | SAE2 | CTIP | SL | SL |

### Functional deficiency of FEN1 induces XPF recruitment to replication forks to mediate 3’ flap processing

Our yeast genetic screening indicated that Rad1 functions as a 3’ flap nuclease for processing 3’ flaps in yeast. We next examined whether XPF, the mammalian homolog of Rad1, plays a similar role in 3’ flap cleavage in mammalian cells. First, we determined if XPF localizes to DNA replication forks. We performed subcellular fractions and chromatin-bound protein isolation in mouse embryonic fibroblast (MEF) cells. In wild-type (WT) cells, XPF was detected in the cytoplasmic, nucleus and chromatin fractions (Figure 1A). However, the nuclear and chromatin-associated levels of XPF were considerably increased in the FEN1-mutant MEFs carrying the F343A/F344A (FFAA) point mutation, which specifically disrupts the interaction between FEN1 and replication core protein PCNA (32), compared with WT cells (Figure 1A). In addition, we found that XPF co-immunoprecipitated with PCNA in 293T cells, and the amount of XPF that was co-IPed with PCNA considerably increased when the cells were treated with LNT-1, a selective FEN1 inhibitor (33) (Figure 1 B). Consistently, co-immunofluorescence (co-IF) staining for XPF and PCNA revealed only a few XPF foci colocalized with PCNA foci in WT MEFs (Figure 1C, 1D and Supplementary Figure S3A, S3B). However, the number of PCNA-colocalized XPF foci was significantly elevated in FFAA or R192Q FEN1 mutant MEF (Figure 1C, 1D) as well as in FEN1 inhibitor-treated MEFs (Supplementary Figure S3A, S3B). Because PCNA foci are widely used as markers of replication forks (34), the observation that XPF co-localized with PCNA foci in FEN1-mutant or FEN1-inhibited cells suggests that FEN1 deficiency induces XPF recruitment to replication forks for 3’ flap processing. To further test this hypothesis, we labeled WT, FFAA and R192Q FEN1-mutant MEFs with EdU as a replication fork marker and performed proximity ligation assay (PLA) to detect replication fork-associated XPF or its interaction partner ERCC1. We observed significantly more XPF-EdU or EdU-ERCC1 PLA foci in the FFAA and R192Q MEFs compared with WT. On average, 10.5±1.0 or 10.3±1.0 XPF-EdU or EdU-ERCC1 PLA foci, respectively, were detected in the FFAA MEFs, and 9.0±1.0 or 13.6±1.2 in the R192Q MEFs (Figure 1E-1G). In contrast, only 2.6 ±0.7 or 3.6 ±0.4 XPF-EdU or EdU-ERCC1 PLA foci, respectively, were observed in WT MEFs (Figure 1E-1G). Consistently, treatment of MEFs or MDA-MB-231 cells with a selective FEN1 inhibitor, LNT-1, significantly increased XPF-EdU or ERCC1-EdU PLA foci (Figure 1H-1J and Supplementary Figure S4A, S4B). These results provide direct evidence that the 3’ flap endonuclease complex XPF-ERCC1 is recruited to replication forks when the FEN1-mediated 5’ flap-based OFM is impaired.

**Figure 1.**
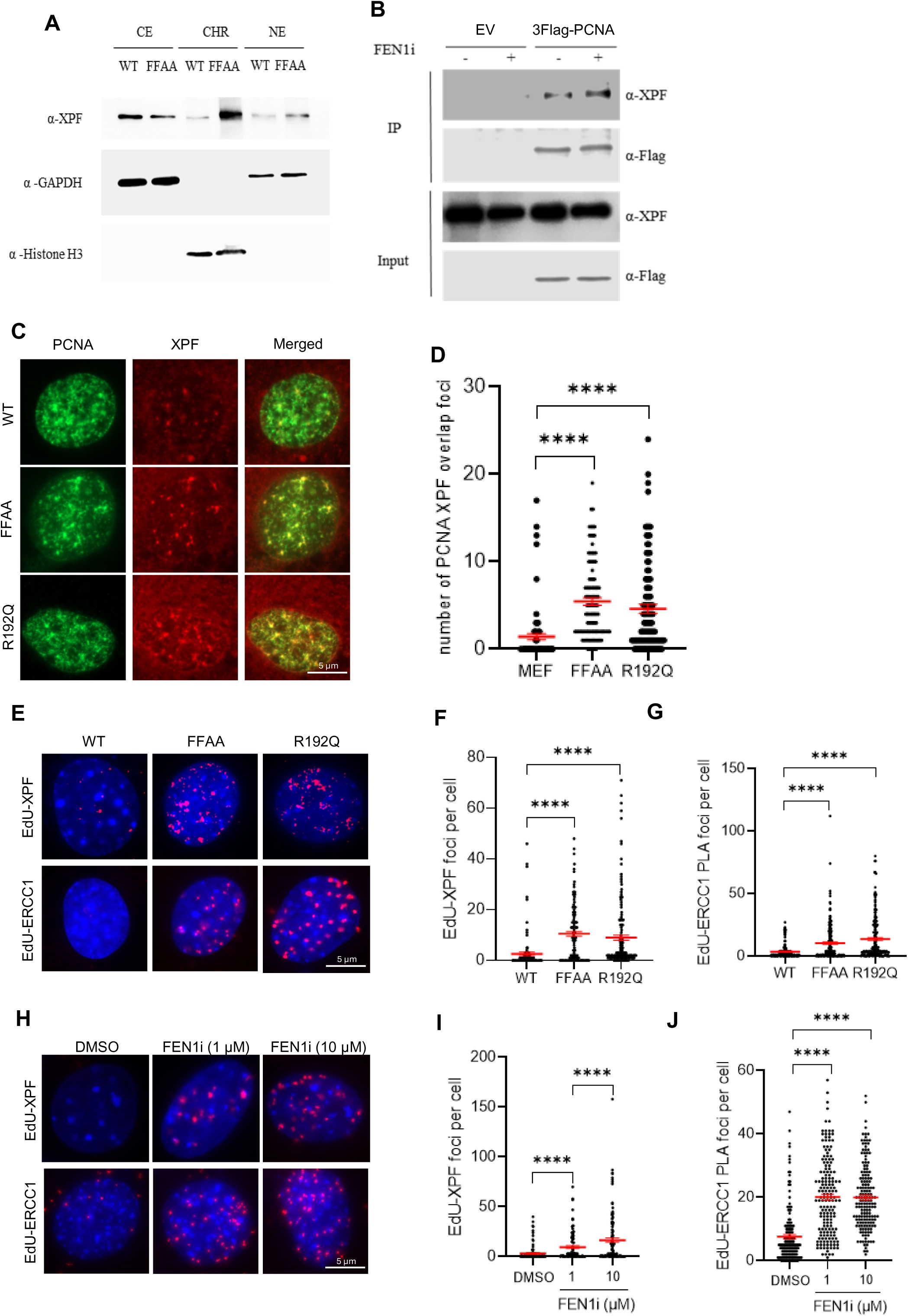
FEN1 functional deficiency FEN1 mutation induces XPF recruitment to replication forks. **(A)** Subcellular fractions and chromatin-bound protein isolation in MEF cells. CE, cytoplasmic. NE, nucleus. CHR, chromatin; **(B)** Immunoblot showing co-IP of Flag-PCNA and XPF in 293T cells. Cells were treated with or without 10 μM FEN1i for 16hr; **(C)** and **(D)** XPF and PCNA co-immunofluorescence (co-IF) staining in WT, FFAA and R192Q mutant MEF cells. (C) Representative microscope images of XPF-PCNA co-IF staining in WT, FFAA and R192Q mutant MEF cells; (D) Quantification of the XPF-PCNA overlap foci number per cell in XPF-PCNA co-IF staining. Data represent mean ± SEM from ≥100 cells per condition. ****P < 0.0001. P value from Student t-test; **(E-G)** PLA performed in WT, FFAA and R192Q mutant MEF cells. (E) Representative microscope images of EdU-XPF and EdU-ERCC1 in WT, FFAA and R192Q mutant MEF cells; Quantification of the EdU-XPF (F) and EdU-ERCC1 (G) PLA foci number in WT, FFAA and R192Q mutant MEF cells. Cells were labeled with10 μM EdU for 20 minutes before harvest. Data represent mean ± SEM from ≥100 cells per condition. ****P < 0.0001. P value from Student t-test; **(H-J)** PLA performed in MEF cells treated with DMSO or FEN1i. (H) Representative microscope images of EdU-XPF and EdU-ERCC1 in MEF cells treated with DMSO or FEN1i; Quantification of the EdU-XPF (I) and EdU-ERCC1 (J) PLA foci number in MEF cells. Cells were treated with DMSO or FEN1i (1 and 10 μM) for 16hr and labeled with10 μM EdU for 20 minutes before harvest. Data represent mean ± SEM from ≥100 cells per condition. ****P < 0.0001. P values were calculated with the Student t-test.

Next, we sought to determine the extent to which XPF is important for removing 3’ flaps in mammalian cells. To quantify 3’ flap levels at the single-cell level, we developed a rolling circle amplification (RCA)-based in situ 3’ flap labeling protocol (Figure 2A). In this approach, which we call 3’ flap-RCA, we designed 3’ flap probe of a degenerate circular ssDNA (55nt), which contains 6 nt random DNA sequences (red section) for 3’ ssDNA flap of varying DNA sequences to anneal. The 3’ flap that annealed to the circular DNA probe was extended by the phi29 polymerase in a rolling circle amplification manner. The extended 3’ flap products are then visualized by hybridizing the ssDNA with a FAM-labeled oligonucleotide (the green fragment) complementary to the ssDNA sequence. We detected green fluorescence signals in FFAA-mutant cells incubated with the circular probe, but not in cells without probe incubation (negative control) (Figure 2B). To further confirm that the green fluorescence signal specifically corresponds to 3’ flaps, we pretreated another slide with *E. coli* EXO1, a 3’ ssDNA exonuclease that removes the 3’ flaps. Pretreatment with *E. coli* EXO1 markedly diminished the green fluorescence signal (Figure 2B), demonstrating that the green fluorescence signal specifically represents nuclear 3’ flaps. Using the 3’ flap-RCA assay, we observed that WT MEFs displayed low 3’ flap signal intensity (Figure 2C, 2D), whereas treatment with NSC143099, a selective XPF inhibitor (XPFi) (35), significantly increased 3’ flap intensity (Figure 2C, 2D). Similarly, the 3’ flap signal intensity in FFAA mutant cells was significantly higher than in WT cells, and treatment of FFAA MEFs with XPFi further enhanced the 3’ flap signal (Figure 2C, 2D). Consistently, XPFi increased 3’ flap signal in HCC827 and MDA-MB-231 cells, two commonly used human cancer cell lines, and FEN1i caused even stronger 3’ flap signals compared with WT cells (Figure 2E, 2F). To further confirm that XPF mediates 3’ flap processing, we generated *Ercc4^−/−^*(XPF knock out) HCC827 and MDA-MB-231 cells. XPF knockout led to a marked accumulation of 3’ flap signals particularly when the cells were treated with FEN1i (Figure 2E, 2F).

**Figure 2.**
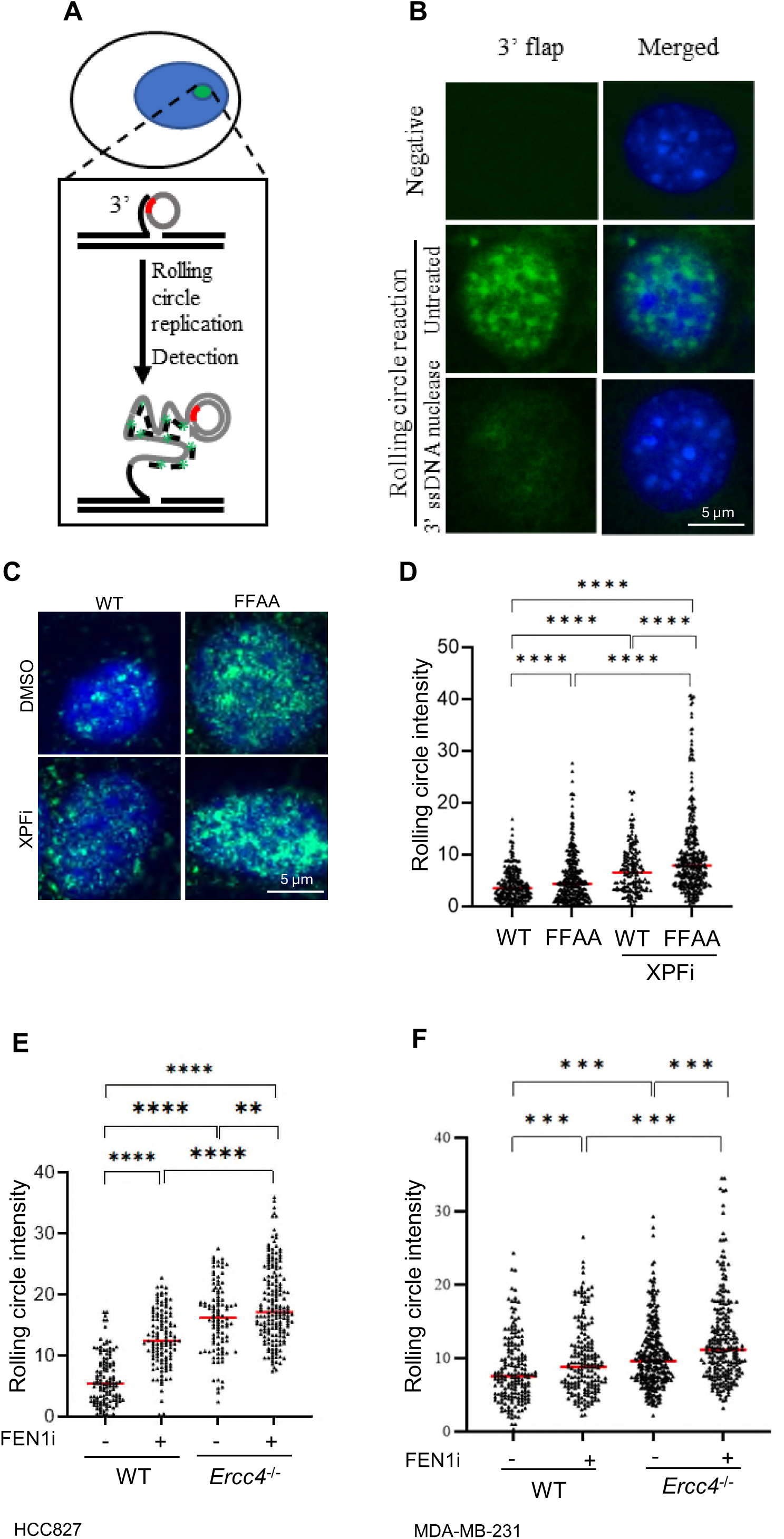
XPF deficient cells accumulate 3’ flaps. **(A)** Diagram of *in situ* detection of 3’flaps using a rolling circle amplification (RCA)-based method; **(B)** Representative microscope images of 3’ flaps detected using RCA in FEN1 FFAA mutant MEF cells, which were fixed using methanol and untreated or treated with 3’ ssDNA nuclease (E. *coli* EXO I) in vitro; **(C)** and **(D)** WT and FFAA mutant MEF cells in RCA. Panel (C) shows representative microscope images of WT and FFAA mutant MEF cells treat with or without XPFi; and panel (D) shows the quantification of RCA in WT and FFAA mutant MEF cells, cells were treated with DMSO or 10 μM XPFi for 16hr. Data represents mean from ≥100 cells per condition. ****P < 0.0001. P value from Student t-test; Quantification of RCA in HCC827 **(E)** and MDA-MB-231 **(F)** WT and *Ercc4^−/−^*cells treated with DMSO or 10 μM FEN1i for 16hr. Data represents 100 cells per condition. **P < 0.01; ***P < 0.001, ***P < 0.0001. P values were calculated with the Student t-test.

### Unremoved 3’flap due to XPF deficiency or inhibition increases replicative DNA damage

Unremoved 5’ flaps and 3’ flaps within OFs prevent OF ligation, leading to the formation of single-strand breaks (SSBs) and double strand breaks (DSBs). To investigate whether deficiency or inhibition of FEN1 or/and XPF causes the accumulation of replicative SSBs, we performed a BrdU alkaline comet assay. In this assay, intact BrdU-labeled nascent DNA forms a compact head, whereas DNA containing single-strand breaks forms a comet-like tail (36,37). In untreated MDA-MB-231 cells, approximately 40% of the BrdU-labeled DNA show no obvious tail (tail length <10 μm), indicating that few SSBs occur in nascent DNA under normal conditions. The mean tail length was 45.2±5.0 μm (Figure 3 A, 3B). However, the tail length of BrdU labeled DNA, representing nascent SSBs, increased dramatically in cells treated with FEN1i or XPFi (Figure 3 A,3B). Moreover, combined treatment both XPFi and FEN1i further increased in the mean tail length to 105.7± 4.2 µm (Figure 3A, 3B). Consistently, *Ercc4^−/−^*(XPF knockout) cells displayed significantly longer comet tails than WT cells, and FEN1i further extended the comet tails in *Ercc4^−/−^*compared with WT (Figure 3C,3D).

**Figure 3.**
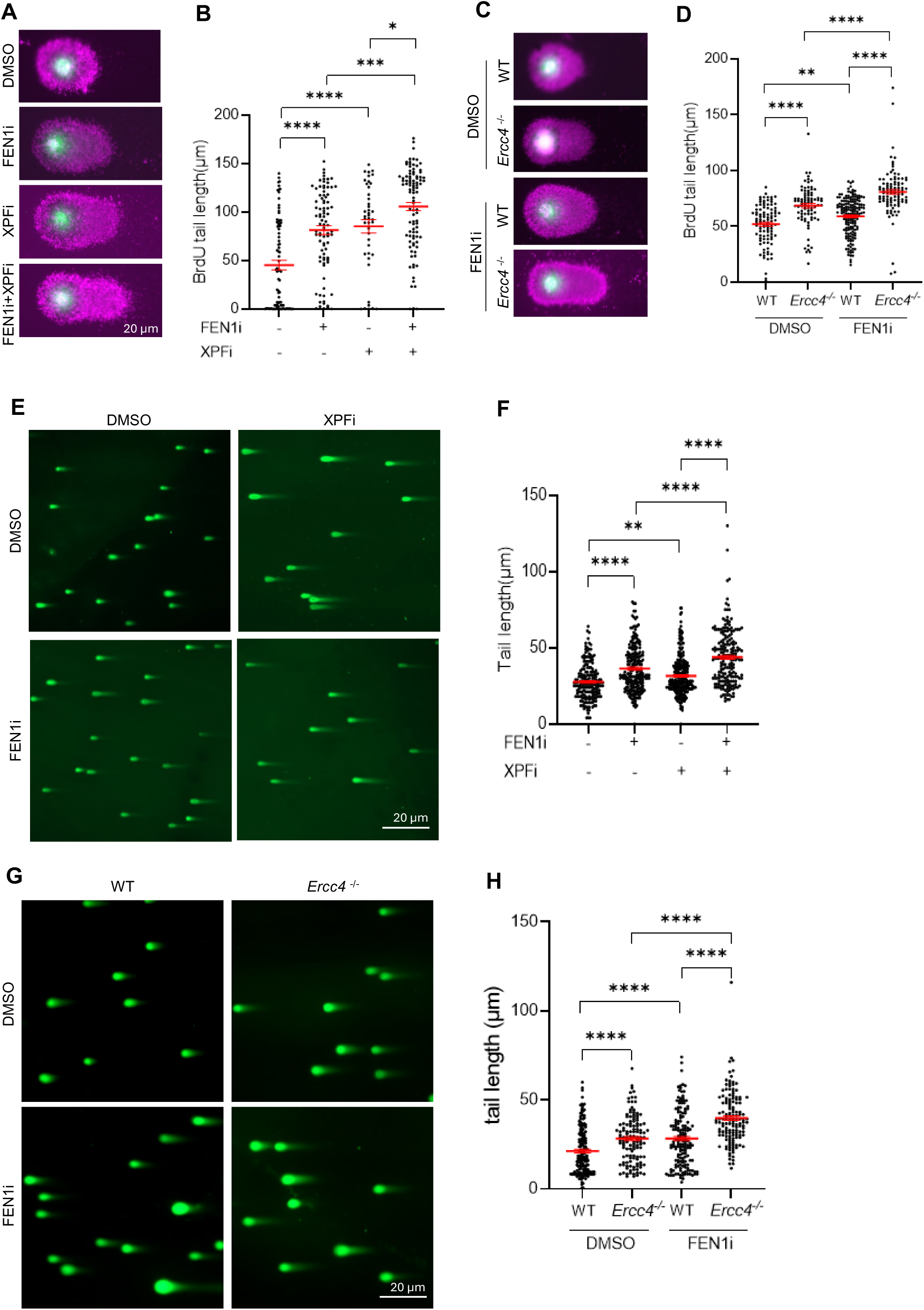
Unremoved 3’flaps due to XPF deficiency or inhibition increase replicative DNA damage. **(A)** and **(B)** BrdU comet assay performed in MDA-MB-231 cells. Panel (A) shows representative microscope images of BrdU COMET assay in MDA-MB-231 cells treated with DMSO/ FEN1i/ XPFi. Panel (B) shows quantification of BrdU tail length per cell in BrdU comet assay. Cells were treated with DMSO/ FEN1i (10 μM)/ XPFi (10 μM) for 16hr and labeled with 20 μM BrdU for 20min before harvest. Data represents mean ± SEM from ≥50 cells per condition. *P<0.05, **P < 0.01, ***P < 0.001, ****P < 0.0001. P values from Student t-test; **(C)** and **(D)** BrdU Comet assay performed in WT and *Ercc4^−/−^*MDA-MB-231 cells. Panel (C) shows representative microscope images of neutral Comet assay in WT and *Ercc4^−/−^* MDA-MB-231 cells treated with DMSO or FEN1i. Panel (D) shows quantification of BrdU tail length per cell in WT and *Ercc4^−/−^*MDA-MB-231 cells. Cells were treated with 10 μM FEN1i and labeled with 20 μM BrdU for 20min before harvest. Data represents mean ± SEM from ≥50 cells per condition. **P < 0.01, ****P < 0.0001. P values from Student t-test; **(E)** and **(F)** Neutral Comet assay performed in MDA-MB-231 cells. Panel (E) shows representative microscope images of neutral Comet assay in MDA-MB-231 cells treated with DMSO/ FEN1i/ XPFi, and panel (F) is quantification of DNA tail length per cell in neutral Comet assay. Cells were treated with DMSO/ FEN1i (10 μM)/ XPFi (10 μM) for 16hr. Data represents mean ± SEM from ≥100 cells per condition. **P < 0.01, ****P < 0.0001. P values from Student t-test; **(G)** and **(H)** Neutral Comet assay performed in WT and *Ercc4^−/−^* MDA-MB-231 cells. Panel (G) shows representative microscope images of neutral Comet assay in WT and *Ercc4^−/−^* MDA-MB-231 cells treated with DMSO or FEN1i. Panel (H) shows quantification of DNA tail length per cell in neutral Comet assay. Cells were treated with DMSO or 10 μM FEN1i for 16hr. Data represents mean ± SEM from ≥100 cells per condition. ****P < 0.0001. P values were calculated with the Student t-test.

Next, we performed a neutral comet assay, which measures the comet tail length corresponding to the level of DSBs in the nucleus. The mean tail length in the MDA-MB-231 cells cultured under normal conditions was 27.6±0.7 μm. This length significantly increased to 36.3±1.0 μm and 31.50±0.8 μm in cells treated with the FEN1i (10 µM, 24 h) or the XPFi (10 µM, 24 h), respectively (Figure 3E, 3F). Combined treatment with both FEN1i and XPFi further increased the mean tail length to 43.8±1.3 μm (Figure 3E, 3F). Consistently, the mean tail length in *Ercc4^−/−^* MDA-MB-231 cells was significantly longer than that in WT cells. Moreover, FEN1i caused a greater increase in DSBs, as reflected by comet tail length, in *Ercc4^−/−^* cells compared with WT cells (Figure 3G, 3H). These results demonstrate that XPF functional deficiency, either through gene deletion or chemical inhibition, like FEN1 inhibition, causes replicative SSBs and consequently DSBs. Furthermore, XPF inhibition shows a synergistic effect with FEN1 inhibition inducing SSBs and DSBs in the genome. These findings suggest that XPF-mediated 3’ flap processing is essential for repairing OFM-related SSBs in human cancer cells, even when FEN1 is functional, and becomes more crucial in FEN1 inhibited or deficient human cancer cells.

### Functional deficiency of XPF or FEN1 invokes distinct DNA damage responses

Accumulation of SSBs and/or DSBs in the genome can activate DNA damage response. Therefore, we analyzed several markers of DSB sensing and repair: γH2AX foci, a marker for histone sensing DSBs (38); RAD51 foci, a marker for activation of homology-directed repair (HDR) of DSBs in S or G2 phase cells (39); and 53BP1 foci, a marker for non-homology-end-joining (NHEJ)-mediated DSB repair (40). We detected significantly higher level of γH2AX foci in FEN1i-treated (13.4±1.1 foci/nucleus) or XPFi-treated (18.0±1.1 foci/nucleus) MDA-MB 231 cells compared with untreated controls (5.2±0.6 foci/nucleus). Notably, combined treatment with XPFi and FEN1i showed a synergistic effect, producing 29.6±1.2 foci per nucleus (Figure 4A, 4B). Interestingly, XPFi treatment caused a marked and statistically significant increase in RAD51 foci, whereas FEN1i alone did not induce RAD51 foci (Figure 4A, 4C). Combined FEN1i and XPFi treatment further enhanced RAD51 foci compared with XPFi alone (Figure 4A, 4C). On the other hand, treatment with either XPFi or FEN1i alone significantly increased in 53BP1 foci, which colocalized with γH2AX, and the combination of both inhibitors exhibited a synergistic effect in inducing 53BP1 foci (Figure 4D-4F). Consistent with the chemical inhibition results, co-IF and western blot analyses show that XPF knockout induced significantly more γH2AX foci than WT cells, especially in the presence of FEN1i (Figure 4G-4I). Together, these results suggest that functional deficiency of FEN1 or XPF activates distinct DNA damage responses. FEN1 deficiency primarily triggers the γH2AX-53BP1 axis, but XPFi activates both the γH2AX-53BP1 axis and the γH2AX-RAD51 axis.

**Figure 4.**
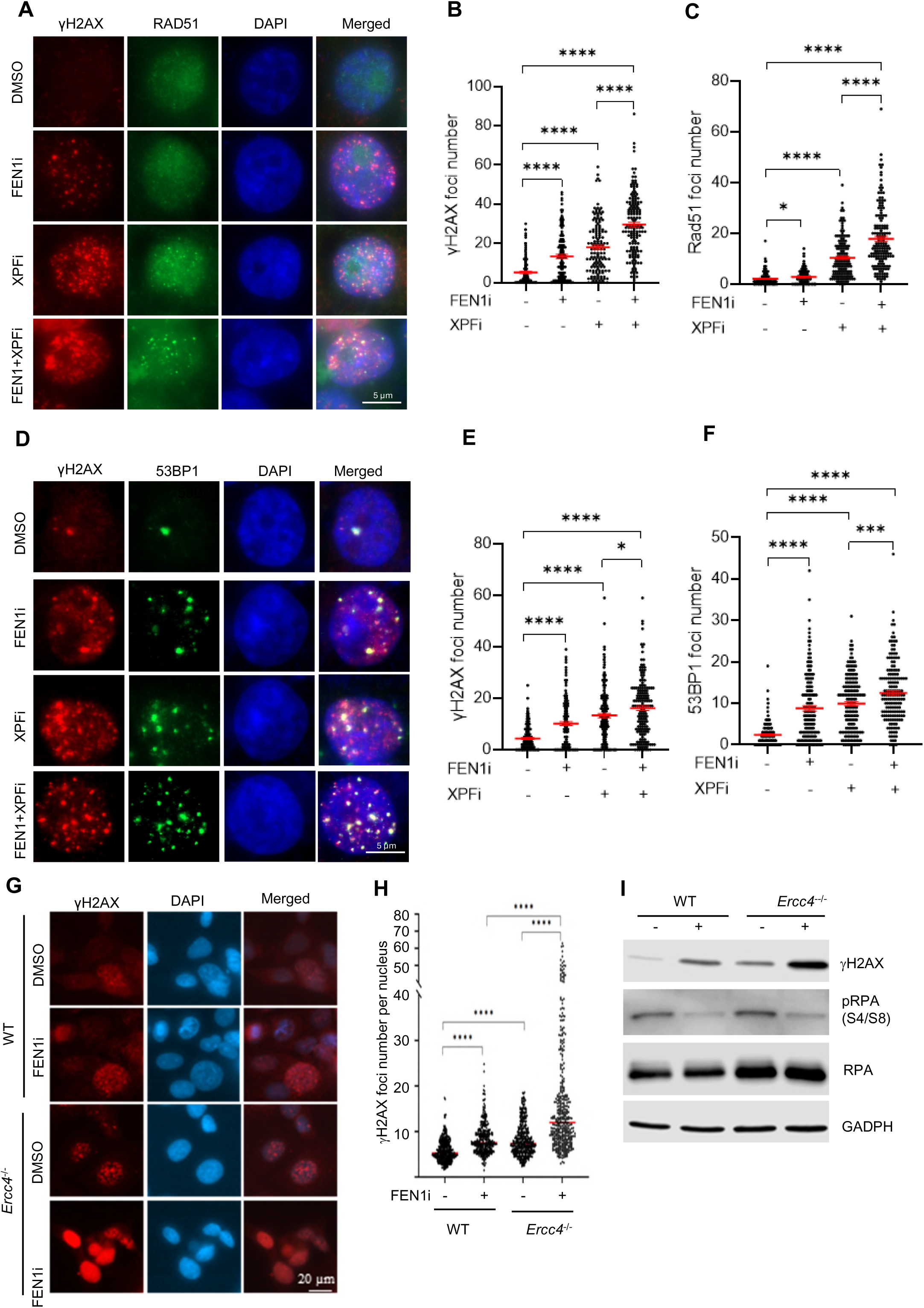
Functional deficiency of XPF or FEN1 invokes distinct DNA damage responses. **(A-C)** γH2AX and RAD51 co-immunofluorescence (co-IF) staining in MDA-MB-231cells. (A) Representative microscope images of γH2AX and RAD51 IF in WT and *Ercc4^−/−^*MDA-MB-231 cells treated with DMSO/ FEN1i/ XPFi. Quantification of γH2AX (B) and RAD51(C) foci number per cell in IF staining. MDA-MB-231 cells were treated with DMSO/ FEN1i (10 μM)/ XPFi (10 μM) for 16hr. Data represents mean ± SEM from ≥100 cells per condition. *P<0.05, ****P < 0.0001. P values from Student t-test; **(D-F)** γH2AX and 53BP1 co-IF staining in MDA-MB-231cells. (D) Representative microscope images of γH2AX and 53BP1 IF in MDA-MB-231 cells treated with DMSO/ FEN1i/ XPFi. Quantification of γH2AX (E) and 53BP1(F) foci number per cell in IF staining. MDA-MB-231 cells were treated with DMSO/ FEN1i (10 μM)/ XPFi (10 μM) for 16hr. Data represents mean ± SEM from ≥100 cells per condition. *P<0.05, ***P<0.001, ****P < 0.0001. P values from Student t-test; **(G)** and **(H)** γH2AX IF staining in WT and *Ercc4^−/−^* MDA-MB-231 cells. (G) Representative microscope images of γH2AX and 53BP1 IF in WT and *Ercc4^−/−^* MDA-MB-231 cells. (H) Quantification of 53BP1 foci number per cell in IF staining. Cells were treated with DMSO or 10 μM FEN1i for 16hr. Data represents median from ≥100 cells per condition. ****P < 0.0001. P values from Student t-test; **(I)** Western blot analysis of γH2AX level in WT and *Ercc4^−/−^* MDA-MB-231 cells. Cells were treated with DMSO or 10 μM FEN1i for 16hr.

### XPF functional deficiency shows a synergy with FEN1 inhibition in causing cell death and DNA mutations

To determine whether XPF is important for cell survival, particularly in cells of functional deficiency in FEN1, we assessed the growth of WT and FFAA MEFs in the presence of varying concentrations of XPFi. We found that FEN1i and XPFi acted synergistically to reduce the viability of HCC827 cells (Figure 5A, 5B). Consistently, FFAA MEFs were significantly more sensitive to XPFi treatment than WT MEFs (Figure 5C). Likewise, *Ercc4^−/−^*MDA-MB-231 cells were markedly more sensitive to FEN1i than WT cells (Figure 5D, Supplementary Figure S5). These findings indicate that functional deficiencies of FEN1 and XPF synergistically suppress cell survival and proliferation, suggesting that XPF completements the role of FEN1 in DNA replication.

**Figure 5.**
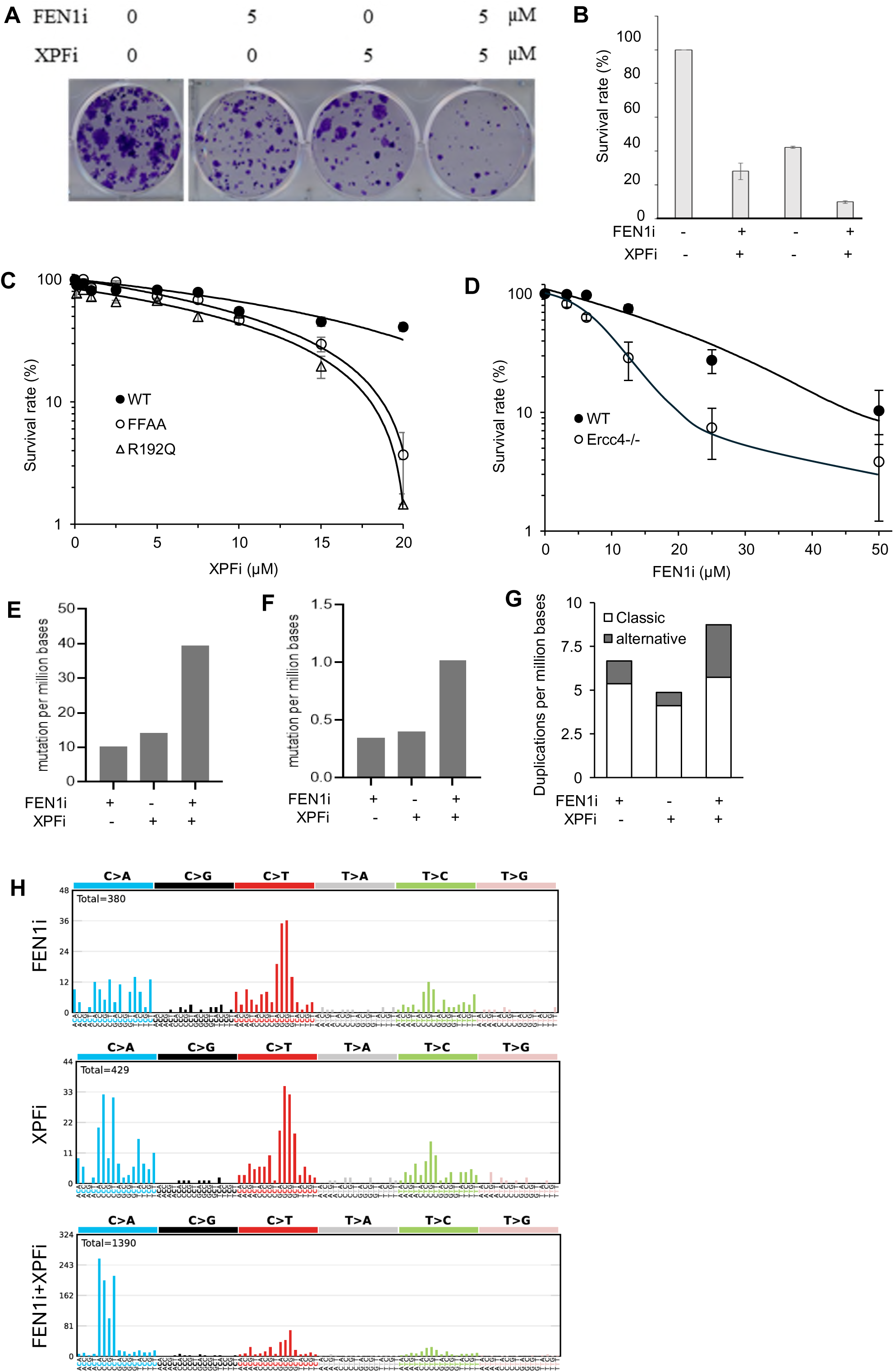
XPF functional deficiency has a synergy with FEN1 inhibition in causing cell death and DNA mutations. **(A) and (B)** Synergistic effects of FEN1i and XPFi in killing HCC827 cells. Panel (A) is the images of clonogenic assay of HCC827 cells in the absence or presence of XPFi or FENi or both. Panel (B) is the quantification of survival rate of the cells under different treatments. The survival rate was untreated control was set as 100%; **(C)** Sensitivity of WT or FEN1 FFAA MEFs to XPFi. Cells were treated with varying concentrations of XPFi for 4 days, and the viable cells were counted. The survival rate of each treatment was calculated relative to untreated control. Data represents SD, n = 3 independent treatments; **(D)** Sensitivity of WT or *Ercc4^−/−^* MDA-MB-231 cells to FEN1i. Cells were treated with varying concentrations of FEN1i for 4 days, and the viable cells were counted. The survival rate of each treatment was calculated relative to untreated control. Data represents mean ± SD, n = 3 independent treatments. **(E-G)** Frequency of single nucleotide variation (SNV) (panel E), small insertion/deletion (Indel) (panel F), and classic or alternative duplications (panel G) as defined by WES. **(H)** Mutation signatures in MDA-MB-231 cells treated with DMSO (untreated control), FEN1i, XPFi, or FEN1i plus XPFi. Occurrence of base substitutions was categorized based on C>A, C>G, C>T, T>A, T>C and T>G and two flanking bases 5′ and 3′ to the mutated base.

Next, we sought to define the frequency and spectra of mutations in MDA-MB-231 cells untreated or treated with the XPFi, the FEN1i, or a combination of both. We isolated the genomic DNA from cells subjected to different treatments and performed whole-exome sequencing (WES). FEN1i treatment caused somatic mutations including 10.2 SNVs, 0.35 small Indels, and 6.7 duplications per million base pairs, whereas XPFi treatment caused somatic mutations including 14.32 SNVs, 0.40 small Indels, and 4.9 duplications per million base pairs (Figure 5E-5G). On the other hand, combined treatment with FEN1i and XPFi showed a synergistic effect, inducing 39.5 SNV, 1.02 small Indels, and 8.7 duplications per million base pairs (Figure 5E-5G). We noted that XPFi on its own caused little alternative duplications, but a combination of XPFi and FEN1i resulted in a remarkable increase in alternative duplications, compared to XPFi or FEN1i alone (Figure 5G). We next performed mutational signature analysis to determine whether FEN1i, XPFi, or their combination generates distinct mutation spectra. In FEN1i-treated cells, C>T mutations were the most predominant SNVs, albeit considerable C>A or T>C mutations were also detected (Figure 5H). In addition, C>T mutations mostly occurred when the “G” nucleotide was upstream of the “C” nucleotide (Figure 5H). This mutation pattern resembles signature SBS15, which is associated with defective mismatch repair (MMR) in human cancers. In XPFi-treated cells, C>A and C>T were the most frequent SNVs (Figure 5H). The C>A mutations mostly occurred when a “C” nucleotide was upstream of the “C” nucleotide, and the C>T mutations mostly occurred when the “G” nucleotide was upstream of the “C” nucleotide (Figure 5H). This pattern is similar to mutation signature SBS44, which is also linked to MMR deficiency in human cancers. Interestingly, cells treated with both FEN1i and XPFi exhibited a strong bias toward the C>A mutations, though considerable C>T mutations were also detected (Figure 5H). This pattern closely resembled mutation signature SBS20, which is found in human cancers harboring POLD1 mutations. Together, these data suggest that the mutagenic processes induced by XPF inhibition are mechanistically related to mismatch repair pathway involved in correcting errors generated by DNA polymerase δ (Pol δ).

## Discussion

Our current studies identify key helicases and 3’ flap endonuclease involved in 3’ flap-based OFM, a newly discovered process that is critical for cell survival under stress conditions. It is well established that the RNA-DNA primer synthesized by Pol α in each Okazaki fragment (OF) must be removed to join OFs into an continuous lagging-strand DNA (10). This process is essential for maintaining genome stability and ensuring cell viability. Under normal physiological conditions, OFM primarily relies on FEN1-mediated cleavage of the RNA-DNA 5’ flap structure generated during Pol δ -mediated strand displacement DNA synthesis. Recently, we discovered that, under replication stress caused by gene deficiencies or environmental stresses that impair 5’ flap processing, eukaryotic cells can activate DUN1 signaling in yeast or the CHK1/2 signaling in mammalian cells to convert unprocessed 5’ flaps into 3’ flaps, which can subsequently be removed by a 3’ flap nuclease (11). This alternative OFM pathway supports cell survival but potentially mutagenic, leading to alternative duplications and base substitutions (41). However, the core enzymes responsible for 3’ flap formation and processing have remained largely unknown. Using yeast genetics analyses, we identified helicases PIF1 and SGS1 and 3’ flap nucleases RAD1 and MUS81 as factors involved in 3’ flap-based OFM. Deletion of either of these genes causes SL with *rad27Δ*, whereas a Pol δ internal tandem duplication (ITD), which limits 5’ flap displacement, rescues the SL phenotype. These observations suggest that SL phenotype depends on existence of the flap structure in the genome and imply that PIF1 or SGS1 may facilitate the conversion of 5’ flaps into 3’ flaps, while RAD1 or MUS81 are likely responsible for 3’ flap cleavage during OFM.

Our studies further define XPF, the human homolog of RAD1, plays a crucial role in removing 3’ flaps during OFM. In mammalian cells, the XPF-ERCC1 complex is generally not localized to replication forks. XPF-PCNA co-IF staining or the XPF-EdU or ERCC1-EdU PLA assay revealed that most MEF cells showed no detectable XPF foci colocalized with PCNA, or any XPF-EdU or ERCC1-EdU PLA signals. However, FEN1 mutation or chemical inhibition dramatically induced XPF localization to replication forks, as demonstrated by both co-IF staining PLA analyses. Intriguingly, a considerable fraction of MDA-MB-231 cells with wild-type FEN1, when cultured under normal conditions, also exhibited recruitment of XPF to replication forks. This suggests that endogenous replication stress or spontaneous DNA damage in human cancer cells activates DDR checkpoints, promoting the formation of 3’ flaps and subsequent recruitment of XPF to replication forks for 3’ flap processing, even in the presence of proficient FEN1 function. Nevertheless, like FFAA MEFs, FEN1i treated MDA-MB-231 cells displayed significantly increased XPF recruitment to replication forks compared with untreated WT cells. Consistently, XPF inhibition or knockout resulted in the accumulation of 3’ flaps and SSBs on newly synthesized DNA, especially in FEN1 deficient MEFs or FEN1 inhibited human cancer cells such as HCC827 or MDA-MB-231.

XPF was originally identified as a core enzyme in nucleotide excision repair (27,28,42,43). It is non-essential for survival of normal cells. Knockout of the yeast XPF homolog RAD1 does not affect yeast cell growth (44). Similarly, XPF knockout mouse embryonic stem cells are viable, and XPF knockout mice can survive until the postnatal stage (45). These observations are consistent with the hypothesis that under normal physiological conditions, FEN1-mediated 5’ flap cleavage serves as the primary mechanism for RNA-DNA primer removal during OFM in mammalian cells. However, human cancer cells experience high levels of endogenous replication stress (3,46), which activate DDR pathways that may promote the conversion of 5’ flaps into 3’ flaps during OFM. Consequently, human cancer cells but not normal cells depend on XPF or other 3’ flap nuclease for efficient OFM and cell survival. Supporting this notion, we observed that HCC827 or MDA-MB-231 cancer cells were more sensitive to XPFi than MEFs. Because of this cancer cell specificity, XPF represents a potential therapeutic target for treating human cancers that have a high possibility to induce 3’ flaps during OFM. Furthermore, in the presence of FEN1i, which blocks the 5’ flap-based OFM, cancer cells become increasingly reliant on the 3’ flap-based OFM, rendering them more sensitive to XPFi.

It is important to note that we observed that XPFi also leads to mutagenesis. We previously showed that unprocessed 3’ flaps can form hairpin structures or anneal to nearby DNA sequences. In such cases, the 3’ flap may undergo self-extension, and subsequent ligation of the extended 3’ flap with the downstream Okazaki fragment can generate alternative duplications. In addition, long 3’ flaps can anneal to a homologous DNA sequence on the sister chromatid, leading to template switching and thus Sister Chromatin Exchange (SCE). Although SCE is generally considered error-free, unequal SCE can result in gene deletions in one chromatid and duplications the other, and have been associated with the development of drug resistance (47). Therefore, efficient removal of 3’ flaps by XPF is critical for maintaining genome stability. XPFi could potentially promote duplications, especially when combined with FEN1i as observed in this study. Given that genetic alterations such as internal tandem duplications, large-scale gene duplications, and chromosome rearrangements can drive drug resistance, therapeutic strategies targeting XPF or other OFM enzymes must take such mutagenic consequences into account to achieve long term beneficial effects. One possible approach to resolve this issue is to block the 3’ flap annealing to nearby sequences via inhibition of RAD52, which mediates microhomology sequence (MHS)-mediated strand annealing (48,49).

## Materials and Methods

### Yeast strains and genetic cross

*Saccharomyces cerevisiae* yeast strain RDKY2672, (MATa, *his3*Δ*200*, *ura3-52*, *leu2*Δ*1*, *trp1*Δ*63*, *ade2*Δ*1*, *ade8*, *hom3-10*, *lys2*Δ*Bgl*) or RDKY2669: *MATα*, *his3*Δ*200*, *ura3-52*, *leu2*Δ*1*, *trp1*Δ*63*, *ade2*Δ*1*, *ade8*, *hom3-10*, *lys2*Δ*Bgl*) was used to create different knockout yeast strains following the published protocol for deletion of a gene from the yeast genome (50). The genotypes of the mutant yeast strains were verified using PCR-based genotyping. The WT and the *rad27*Δ strain (RDKY2608: a, *his3*Δ*200*, *ura3-52*, *leu2*Δ*1*, *trp1*Δ*63*, *ade2*Δ*1*, *ade8*, *hom3-10*, *lys2*Δ*Bgl*, *rad27*Δ*::URA3*) were gifts from Dr. Richard D. Kolodner *rad27Δ::LEU2 or rad27Δ::LEU2 pol3-ITD*::HIS3 mutant yeast strains (MATa or MATα) used in this study were generated in our previous study (11).

Random spore analysis was carried out following a standard protocol (51), in order to verify the SL phenotype of *rad27*Δ with *gene deficiency in a helicase or nuclease as listed in Table 1, and t*o assess if *pol3* ITD rescues the SL phenotype. Briefly, he diploid yeast mutant cells were created by genetic crosses of the *rad27*Δ::LEU2 or *rad27*Δ::LEU2 *pol3* ITD::HIS3 mutant strain with the helicase or nuclease deletion strain (URA3 as the selection marker). The ascospores produced from the diploid mutant cells were isolated as previously described. The haploid was verified by PCR analysis for the status of carrying either the MATa or the MATα allele. The following primers were used: MATa forward primer: 5’ ACTCCACTTCAAGTAAGAGTTTG 3’, MATα forward primer: 5’ GCACGGAATATGGGACTACTTCG 3’, and MAT reverse primer: 5’ AGTCACATCAAGATCGTTTATGG 3’. The spores were cultured in YPD medium, and the viable spores were selected on the SC-Leu-His nutrition deficient medium plates or SC-Leu-His-Ura plates as previously reported (11).

### Cell culture

HCC827, MDA-231, Mouse embryonic fibroblast (MEF) cells were cultured in DMEM medium supplemented with 10% fetal bovine serum (FBS; Gibco) and 1% penicillin-streptomycin at 37 °C. The HCC827 and MDA-231 cell lines were obtained from the ATCC and the MEF cell lines (WT, FFAA, R192Q) were isolated from mouse embryos (E13.5) at Animal Resource Center of COH. All cell lines were subjected to mycoplasma testing twice per month and found to be negative.

### Subcellular fractions and chromatin-bound protein isolation

To isolate cytoplasmic extracts (CEs) and unclear extracts (NEs), cells were collected and suspended in CE buffer (10 mM HEPES-KOH [pH 7.5], 60 mM KCl, 1 mM DTT, 1 mM EDTA, 0.075% Nonidet *P-*40, 1 ×protease inhibitor cocktail) for 10 min on ice. After centrifugation (1000 × *g*, 10 min), the supernatant (CE) was collected. The pellet was then washed with CE buffer without Nonidet *P-*40 twice and resuspended in NE buffer (25% glycerol, 20 mM Tris-HCl, 420 mM NaCl, 1.5 mM MgCl_2_, 0.2 mM EDTA, 1 × protease inhibitor cocktail) for 1 h on ice with intermittent vortex. After centrifugation (20,000 × *g*, 15 min), the supernatant (NE) was collected and subjected to western blot analysis or stored at −80 °C.

To fractionate cytoplasmic, soluble nuclear and chromatin-bound proteins, the cells were lysed in five volumes of ice-cold Buffer A (50 mM HEPES-KOH [pH 7.5], 140 mM NaCl, 1 mM EDTA [pH 8.0], 10% glycerol, 0.5% Nonidet *P-*40, 0.25% Triton X-100, 1 mM DTT, 1 × protease inhibitor cocktail). After centrifugation (700 × *g*, 10 min), the supernatant was collected as the cytoplasm fraction and pellets were washed with buffer A and resuspended in ice-cold Buffer B (10 mM Tris-HCl [pH 8.0], 200 mM NaCl, 1 mM EDTA [pH 8.0], 0.5 mM EGTA [pH 8.0], 1 × protease inhibitor cocktail). After extraction and centrifugation (20,000 × *g*, 10 min), the soluble NE was collected. The pellet was washed with Buffer B and resuspended in Buffer C (10 mM Tris-HCl [pH 8.0], 200 mM NaCl, 1 × protease inhibitor cocktail). The suspension was sonicated and centrifuged (20,000 × *g*, 10 min), and the resulting supernatant (chromatin-bound proteins) was collected. All fractionated samples were subjected to western blot analysis or stored at - 80 °C.

### Immunoprecipitation (IP)

For immunoprecipitation (IP), expression plasmids were transfected into HEK293 cells. HEK293 cells were harvested 64h after transfection, after which the pellet was lysed with Lysis buffer (20mM Tris-HCl [pH 7.5], 150mM NaCl, 10% glycerol, 0.5% NP-40, 10 mM NaF, 1 mM PMSF, 1 μg/mL leupeptin, and 1 μg/mL aprotinin). The lysate was subjected to centrifugation at 40,000*×g* for 15 min, after which the supernatant incubated with anti-FLAG M2 conjugated agarose beads at 4 °C for 4 h. The beads were washed four times with IP buffer (20 mM Tris-HCl [pH 7.5], 150 mM NaCl, 5 mM MgCl2, 10% glycerol, 0.1% NP-40, 1 mM DTT, and 1 mM PMSF) and incubated with 400 μg/mL 3× Flag peptide in IP buffer for 1-2 h. Subsequently, the eluted complexes were analyzed with sodium dodecyl sulfate-polyacrylamide gel electrophoresis (SDS-PAGE).

### Proximity ligation assay (PLA)

Cells were seeded at a density of 2 × 10⁵ cells per well in 24-well plates containing sterile glass coverslips and cultured overnight in DMEM. Cells were treated with the FEN1 inhibitor LNT1 at the concentrations indicated in the figure legends and incubated overnight. Following treatment, cells were fixed with 4% paraformaldehyde for 30 minutes at room temperature, then permeabilized with 0.2% Triton X-100 for 15 minutes. Cells were blocked with Duolink® Blocking Solution (Sigma) for 1 h and incubated overnight at 4°C with primary antibodies diluted in Duolink® Antibody Diluent (Sigma). After washing, cells were incubated with Duolink® In Situ PLA® Probe Anti-Rabbit MINUS and Anti-Mouse PLUS (Sigma) for 1 h at 37°C. PLA reactions were performed using the Duolink® In Situ Detection Reagents Red kit (Sigma), including ligation for 30 minutes at 37°C, followed by rolling circle amplification with polymerase for 100 minutes at 37°C. PLA signals were visualized and recorded using a Zeiss Observer II or LSM 900 confocal microscope.

### Click-iT EdU Cell Proliferation Assays

Cells were seeded at a density of 2 × 10⁵ cells per well in 24-well plates containing sterile glass coverslips and cultured overnight in the presence of the FEN1 inhibitor LNT1, as indicated. To visualize subnuclear localization linked to nascent DNA synthesis, cells were incubated with 10 μM EdU for 20 minutes prior to fixation. Fixation and permeabilization were performed as described in the “Proximity Ligation Assay” section. EdU-labeled DNA was then detected by a Click reaction using a solution containing 2 mM CuSO₄, 10 μM Alexa Fluor 488 azide (or azide-biotin for PLA), and 50 mM sodium ascorbate in PBS, incubated for 1 h at room temperature, protected from light when necessary. Afterward, cells were rinsed with PBS and blocked with Image-iT™ FX Signal Enhancer (Invitrogen) for 30 minutes at room temperature. Primary and secondary antibodies were diluted and applied as described in the Immunofluorescence” section. Images were acquired using a Zeiss Observer II or LSM 900 confocal microscope.

### Immunofluorescence microscopy

The subnuclear localization sites of XPF, PCNA, RAD51, 53BP1, γH2AX were determined using indirect immunofluorescence analysis. Cells were cultured on coverslips to ∼50% confluence. After washing with PBS, cells were fixed with 4% paraformaldehyde for 30 min at room temperature. Next, the cells were permeabilized for 15 min with 0.2% Triton X-100 in PBS buffer, washed three times with 0.05% Tween-20 in PBS and blocked with 5% BSA for 30 min. Next, the cells were incubated with the primary antibodies diluted in 1% BSA/PBS for 90 min. After washing, the cells were incubated with the secondary antibodies for 45 min. After being washed three times, the cells were mounted with ProLong Gold antifade reagent with DAPI (Invitrogen). Immuno-fluorescence images were analyzed and recorded using Zeiss LSM800 confocal microscope or Observer II fluorescence microscope.

### Rolling circle amplification (RCA)-based 3’ flap detection assay

An RCA-based assay was developed to detect 3’ flaps in the nucleus in situ. To probe 3’ flaps in the nucleus, a degenerate circular ssDNA was constructed. The 5’ end phosphorylated DNA oligo (5’-PO4-GTTTAAGCGTCTTAANNNNNNGCGAGACGGACTCGCATTCACTGGAAAGAGAGTAGTACAG CA GCCGTCAAGAGTGTCTAGTTGTGTCATC-3’) was synthesized. The oligo contains a region of 6 nt random DNA sequences (NNNNNN), which is used for 3’ ssDNA flap of varying DNA sequences to anneal to. The oligo was circularized by incubation the oligo with a template oligo (5’ TTAAGACGCTTAAACGATGACAGAAC-3’) at 1:2 ratio and T4 DNA ligase (16°C, 16 h). The circular product was purified using a 15% denaturing PAGE. To detect 3’ flaps in situ, the cells that were cultured on the coverslip were fixed by 4% paraformaldehyde (room temperature, 10 min). The cells were permeabilized with 0.2% triton X100 (Room temperature, 15 min) and blocked with the blocking reagent (SIGMA) (Room temperature, 30 min). After washing with PBS buffer, the coverslip was incubated with the circular probe (100 nM) in PBS buffer (Room temperature, 30 min). The coverslip was extensively washed with PBS buffer to remove the free circular probe. The coverslip was incubated with RCA reaction supermix containing the RNase H to remove that RNA that annealed to the circular probe and the phi29 polymerase (New England Biolabs) in the phi29 polymerase reaction buffer supplied by the manufacture. RCA reactions were carried out at 30°C for 1 hour and stopped by washing with the hybridization buffer containing 20% formamide 20 mM EDTA. The washed coverslip was incubated with the detection oligo (5’-FAM-AGACGGACTCGCATTCACTGGA-3’, 1 µM) in the hybridization buffer. To visualize the extended 3’ flap, the amplified ssDNA sequence is hybridized with a FAM-oligo (the green fragment) that is complementary to the ssDNA sequence, and the green fluorescence signal was visualized under a fluorescence microscope.

### BrdU comet assay

Neutral and BrdU alkaline comet assays were performed using the Comet Assay Kit (Trevigen, 4250-050). For the BrdU alkaline comet assay, cells were incubated with 20 μM BrdU. Cells were harvested and resuspended in PBS as a concentration of approximately 1X10^5^ cells/ mL. A volume of 5 µL of the cell suspension was mixed with 50 µL of 0.5% low-melting-point agarose (LMPA) at 37 °C and immediately layered onto a slide. The slides were placed at 4 °C for 10 minutes to solidify. After agarose solidification, the slides were immersed in cold lysis buffer (Trevigen, 4250-050) for at least 1 h at 4 °C. Following lysis, the slides were immersed in Alkaline Unwinding Solution (200 mM NaOH, 1 mM EDTA) to unwind the DNA. Electrophoresis was carried out at 21 V for 30 minutes at 4 °C in Alkaline Electrophoresis Solution (200 mM NaOH, 1 mM EDTA). After electrophoresis, slides were gently washed with distilled water, fixed in 70% ethanol for 10 minutes, and allowed to air dry in 37 °C. Then slides were stained with anti-BrdU (BD 347580) antibodies and secondary antibodies. Slides were imaged on a observe II microscope and ZEN 3.1 software. The tail moment of comet was measured by Open Comet of image J.

### Neutral comet assay

Neutral and BrdU alkaline comet assays were performed using the Comet Assay Kit (Trevigen, 4250-050). The neutral comet assay was performed to evaluate DNA double-strand breaks as previously described. Briefly, cells were harvested and resuspended in PBS as a concentration of approximately 1×10^5^ cells/ mL. A volume of 5 µL of the cell suspension was mixed with 50 µL of 0.5% low-melting-point agarose (LMPA) at 37 °C and immediately layered onto a slide. The slides were placed at 4 °C for 10 minutes to solidify. After agarose solidification, the slides were immersed in cold lysis buffer (Trevigen, 4250-050) for at least 1 h at 4 °C. Following lysis, the slides were rinsed with TBE, then placed in an electrophoresis tank filled with the TBE. Electrophoresis was carried out at 21 V for 45 minutes at 4 °C. After electrophoresis, slides were gently washed with distilled water, fixed in 70% ethanol for 10 minutes, and allowed to air dry in 37 °C. The DNA was stained with SYBR Green for 30 minutes in dark. Comet images were captured using a fluorescence microscope, and tail moment were analyzed using Open Comet software in image J.

### Cell survival assay

For the cell survival assay, 50,000 cells were plated into each well of a 12-well plate containing DMEM medium (10% FBS, 1% penicillin-streptomycin). The plates were treated with a range of dose of FEN1 inhibitor LNT1 or XPF inhibitor. After 4 days of incubation at 37 °C, the cells were collected and counted.

### Whole-exome sequencing (WES) and data analysis

WES were conducted following the published protocol as we previously did (14). Genomic DNA isolated from MDA-MB-231 cells treated with FEN1i, XPFi, or in combination. Exons were enriched using all human exon probe set (Agilent) and the WES library was prepared using the KAPA DNA HyperPrep kit (Roche). Sequencing on the WES library was carried out on an Illumina HiSeq2500 using a paired end mode. The quality of sequencing reads was analyzed using FastQC. Trim galore (version 0.6.10) was used to remove any adaptor sequence. The paired end reads with reads were longer than 35 bp from both ends after trimming were subject to further analysis. Bowtie2 (version 2.4.1) was used to map the sequence reads to the human reference genome hg38. The sorted and indexed sequence alignment map (SAM) file was used to create the sorted BAM file using the MarkDuplicates from Picard toolbox (version 2.27.5). Single nucleotide variations or small insertions or deletions were analyzed using Mutect2 (v.4.1.4.0) with the parameter “--panel-of-normals 1000g_pon.hg38.vcf.gz, --germline-resource af-only-gnomad.hg38.vcf.gz, --af-of-alleles-not-in-resource 0.0000025”. Only the SNPs or indels with “PASS” were used to perform further analysis. Tandem duplications and other structural variations were analyzed using Pindel (version 0.2.5b9). Somatic duplications were scored if at least 2 supporting tracks from the upstream and at least 2 supporting tracks from the downstream of the break points in the sample were detected but no supporting tracks were detected in the WT control. The frequencies of the duplication or other mutations were estimated by dividing the number of somatic mutations or structure variations by size of mouse exome (∼30 million). SigProfilerClusters v.1.0.115 was used for clustered mutation analysis. A window size of 1 Mb was used to adjust intramutational distances based on local mutation density, and variant allele frequencies were also taken into account during the subclassification process. The SigProfilerAssignment v.0.2.56 was used to for mutation signature analysis and comparison to the reference signatures derived from Catalogue of Somatic Mutations in Cancer (COSMIC) database (52).

## Data Availability

The WES sequence data have been deposited into the NCBI database. The accession number is PRJNA1354089.

## Supplementary Data

Supplementary Tables and Figures.

## Acknowledgements

We thank Light Microscopy Digital Imaging (LMDI) Shared Resource at City of Hope for assistance with microscopy. Research reported in this publication included work performed in the LMDI Shared Resource supported by the National Cancer Institute of the National Institutes of Health under grant number P30 CA033572. The content is solely the responsibility of the authors and does not necessarily represent the official views of the National Institutes of Health. This work was supported by NIH grants R50 CA211397 to L.Z. and R01 CA073764 and R01 CA279840 to B.S.

## Author contributions

K. Li, G. Shi, YY Wang, M. Zhou conducted biochemical and cellular experiments. F. Yang, Y Lei, YX Wang, and L. Zheng conducted DNA sequence analysis. L. Zheng, K. Li, and B. Shen wrote the manuscript. L. Zheng and B. Shen designed the experiments and supervised the execution of the entire project.

