## Supplementary Figures and Tables for "XPF mediates 3’ flap processing for FEN1-independent Okazaki fragment maturation"

Table S1: Random spore analysis of the SL phenotype of *rad1*Δ *rad27*Δ and suppression of the SL phenotype by *pol3* ITD

| Genetic cross | Colony Number (YPD plates) | Colony Number (SC plates) | Observed ratio | Expected Ratio | Viability |
|---|---|---|---|---|---|
| *rad1*Δ::Trp1 × *rad27*Δ::Ura3 | 103 | *rad1*Δ *rad27*Δ 3 | 0.029 | 0.25 | NO |
| *rad1*Δ::Trp1 × *rad27*Δ::Ura3 *pol3* ITD::His3 | 154 | *rad1*Δ *rad27*Δ *pol3* ITD 17 | 0.110 | 0.125 | YES |
| *rad1*Δ::Trp1 × *pol3* ITD::His3 | 645 | *rad1*Δ *pol3* ITD 241 | 0.374 | 0.25 | YES |

Table S2: Random spore analysis of the SL phenotype of *mre11*Δ *rad27*Δ and suppression of the SL phenotype by *pol3* ITD

| Genetic cross | Colony Number (YPD plates) | Colony Number (SC plates) | Observed ratio | Expected Ratio | Viability |
|---|---|---|---|---|---|
| *mre11*Δ::Trp1 × *rad27*Δ::Ura3 | 961 | *mre11*Δ *rad27*Δ 6 | 0.006 | 0.25 | NO |
| *mre11*Δ::Trp1 × *rad27*Δ::Ura3 *pol3* ITD::His3 | 768 | *mre11*Δ *rad27*Δ *pol3* ITD 3 | 0.004 | 0.125 | NO |
| *mre11*Δ::Trp1 × *pol3* ITD::His3 | 606 | *mre11*Δ *pol3* ITD 174 | 0.287 | 0.25 | YES |

Table S3: Random spore analysis of the SL phenotype of *mus81*Δ *rad27*Δ and suppression of the SL phenotype by *pol3* ITD

| Genetic cross | Colony Number (YPD plates) | Colony Number (SC plates) | Observed ratio | Expected Ratio | Viability |
|---|---|---|---|---|---|
| *mus81*Δ::Trp1 × *rad27*Δ::Ura3 | 81 | *mus81*Δ *rad27*Δ 2 | 0.025 | 0.25 | NO |
| *mus81*Δ::Trp1 × *rad27*Δ::Ura3 *pol3* ITD::His3 | 400 | *mus81*Δ *rad27*Δ *pol3* ITD 61 | 0.153 | 0.125 | YES |
| *mus81*Δ::Trp1 × *pol3* ITD::His3 | 81 | *mus81*Δ *pol3* ITD 18 | 0.222 | 0.25 | YES |

Table S4: Random spore analysis of the SL phenotype of *sae2*Δ *rad27*Δ and suppression of the SL phenotype by *pol3* ITD

| Genetic cross | Colony Number (YPD plates) | Colony Number (SC plates) | Observed ratio | Expected Ratio | Viability |
|---|---|---|---|---|---|
| *sae2*Δ::Trp1<br>×<br>*rad27*Δ::Ura3 | | *sae2*Δ *rad27*Δ | | 0.25 | NO |
| *sae2*Δ::Trp1<br>×<br>*rad27*Δ::Ura3<br>*pol3* ITD::His3 | 530 | *sae2*Δ *rad27*Δ *pol3* ITD<br>3 | 0.006 | 0.125 | NO |
| *sae2*Δ::Trp1<br>×<br>*pol3* ITD::His3 | | *sae2*Δ *pol3* ITD | | 0.25 | |

Table S5: Random spore analysis of the SL phenotype of *sgs1*Δ *rad27*Δ and suppression of the SL phenotype by *pol3* ITD

| Genetic cross | Colony Number (YPD plates) | Colony Number (SC plates) | Observed ratio | Expected Ratio | Viability |
|---|---|---|---|---|---|
| *sgs1*Δ::Trp1<br>×<br>*rad27*Δ::Ura3 | 68 | *sgs1*Δ *rad27*Δ<br>0 | 0 | 0.25 | NO |
| *sgs1*Δ::Trp1<br>×<br>*rad27*Δ::Ura3<br>*pol3* ITD::His3 | 38 | *sgs1*Δ *rad27*Δ *pol3* ITD<br>5 | 0.132 | 0.125 | YES |
| *sgs1*Δ::Trp1<br>×<br>*pol3* ITD::His3 | 68 | *sgs1*Δ *pol3* ITD<br>16 | 0.235 | 0.25 | YES |

Table S6: Random spore analysis of the SL phenotype of *srs2*Δ *rad27*Δ and suppression of the SL phenotype by *pol3* ITD

| Genetic cross | Colony Number (YPD plates) | Colony Number (SC plates) | Observed ratio | Expected Ratio | Viability |
|---|---|---|---|---|---|
| *srs2*Δ::hph × *rad27*Δ::Leu2 | 115 | *srs2*Δ *rad27*Δ 0 | 0 | 0.25 | NO |
| *srs2*Δ::hph × *rad27*Δ::Leu2 *pol3* ITD::His3 | 223 | *srs2*Δ *rad27*Δ *pol3* ITD 7 | 0.031 | 0.125 | NO |
| *srs2*Δ::hph × *pol3* ITD::His3 | 1226 | *srs2*Δ *pol3* ITD 254 | 0.207 | 0.25 | YES |

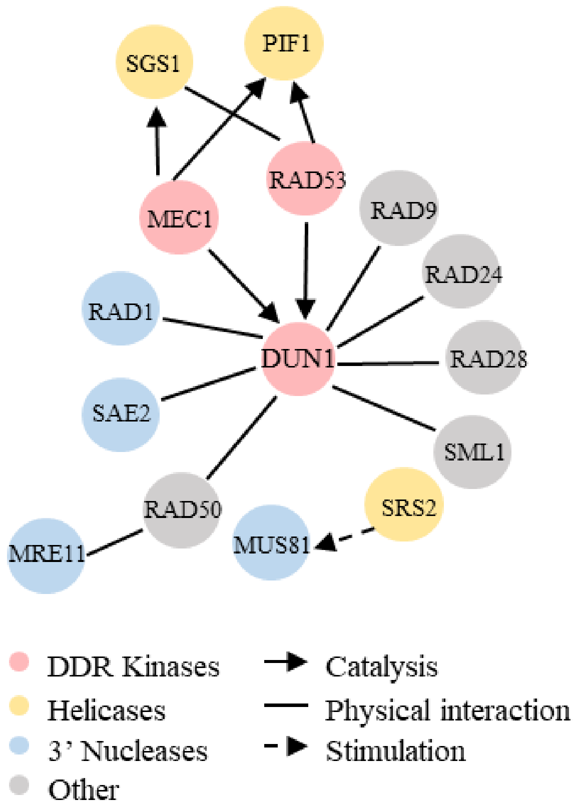


**Supplementary Figure S1**: **Physical interactions of DNA repair proteins with the Mec1-Rad53-Dun1 axis.** Saccharomyces Genome Database (SGD) was surveyed for DUN1 interaction helicases and nucleases.

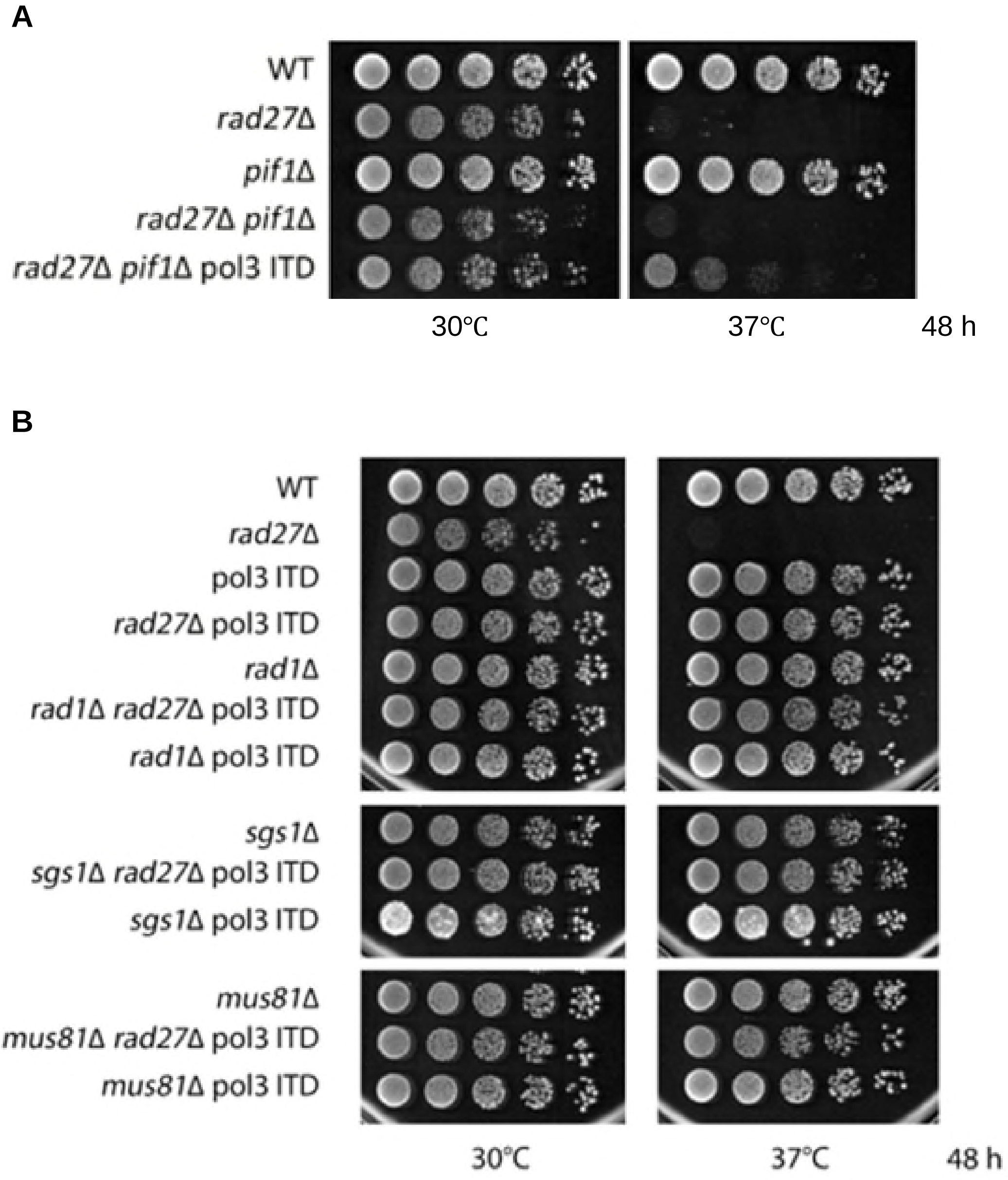


**Supplementary Figure S2: Spot assays to verify the viability of yeast cells. (A)** Spot assays on WT, *rad27*Δ, *pif1Δ*, *rad27*Δ *pif1Δ, or rad27*Δ *pif1Δ* pol3-ITD at 30℃ (optimal temperature) or 37℃ (restrictive temperature). **(B)** Spot assays on WT, *rad27*Δ, *rad1Δ*, *sgs1Δ, mus81Δ, rad27*Δ *rad1Δ, rad27*Δ *sgs1Δ, rad27*Δ *mus811Δ, or rad27*Δ *rad1Δ pol3-ITD, rad27*Δ *sgs1Δ pol3-ITD, rad27*Δ *mus811Δ pol3-ITD* at 30℃ or 37℃.

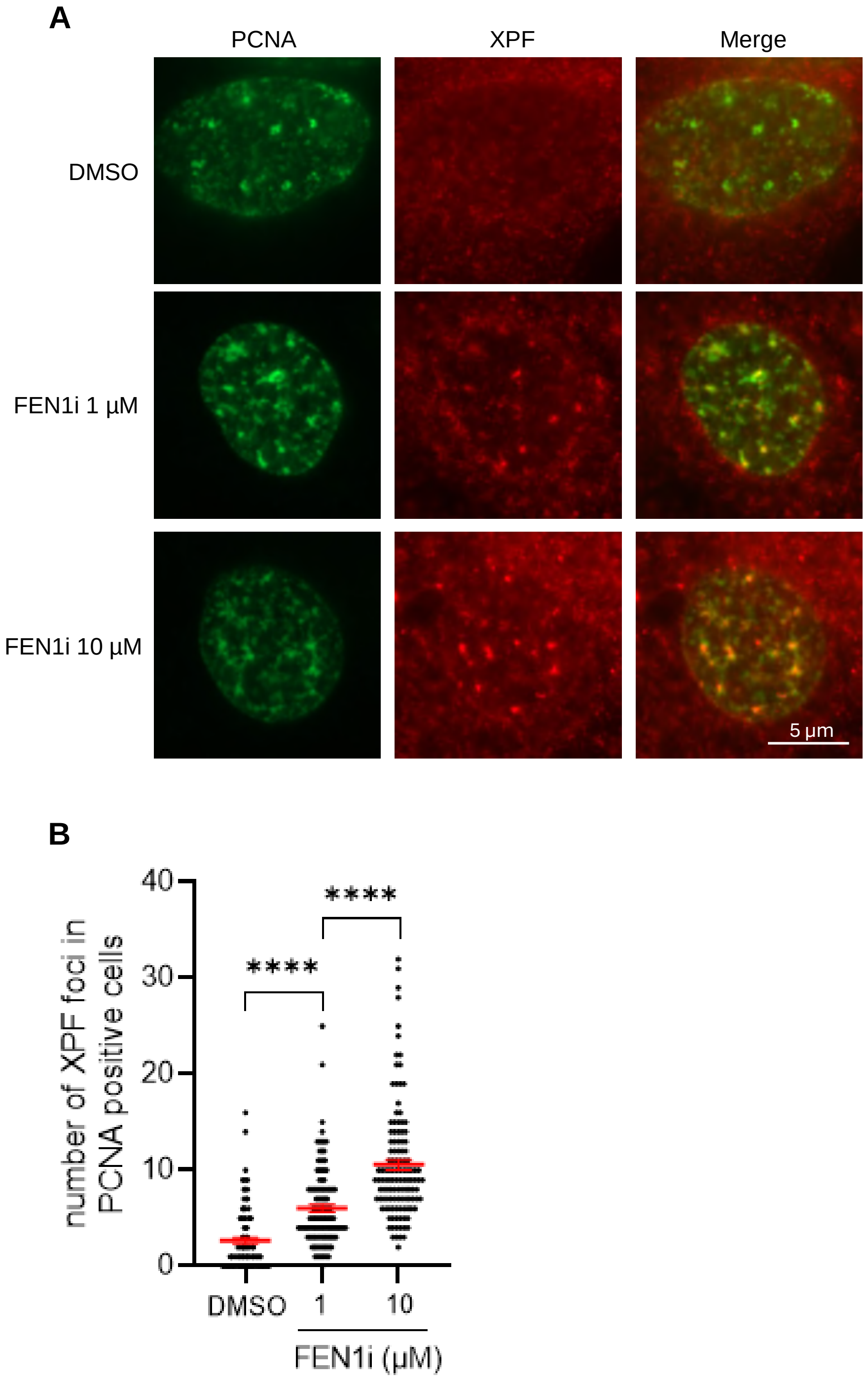


**Supplementary Figure S3: co-IF staining of PCNA and XPF in MEFs in the absence or presence of FEN1i.** MEFs were treated with DMSO or FEN1i (1 µM or 10 µM LNT-1) for 16hr. Co-IF staining of PCNA and XPF was carried out. **(A)** Representative microscope images of XPF-PCNA co-IF staining in MEF cells treated with DMSO or FEN1i; **(B)** Quantification of the XPF-PCNA co-localized foci number per cell. Data represent mean ± SEM from ≥100 cells per condition. **** $p < 0.0001$. P values were calculated with the Student t-test.

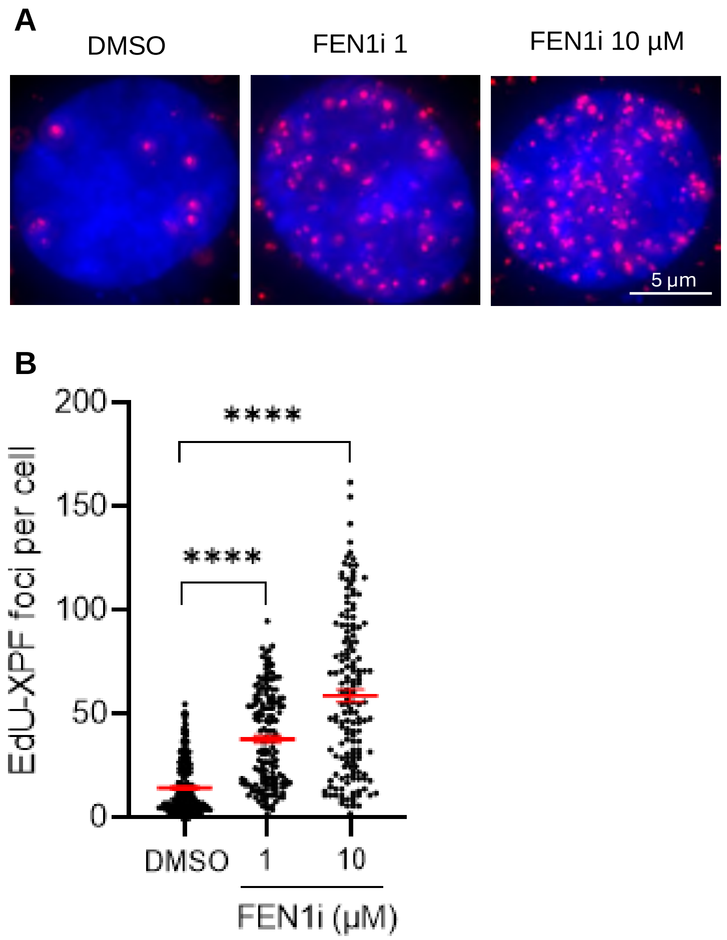


**Supplementary Figure S4: The EdU-XPF PLA assay shows XPF recruitment to nascent DNA** in MDA-MB-231 cells. **(A)** Representative microscope images of PLA performed in MDA-MB-231 cells treated with DMSO or FEN1i; **(B)** Quantification of the EdU-XPF PLA foci number in MDA-MB-231 cells. Cells were treated with DMSO or FEN1i (1and 10 µM) for 16hr and labeled with10 µM EdU before harvest. Data represent mean ± SEM from ≥100 cells per condition. ****P < 0.0001. P value from Student t-test.

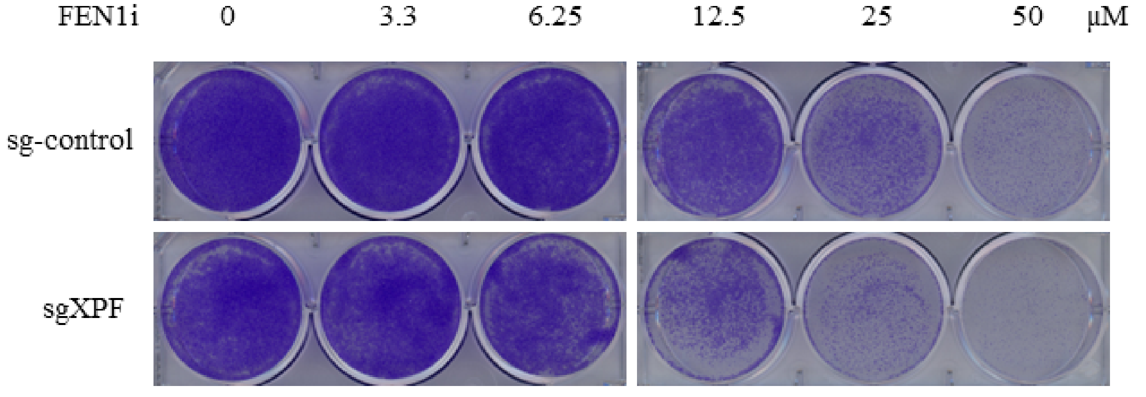


**Supplementary Figure S5: Cell viability assay in WT (sg-control) or ERCC4-/- (sgXPF)** MDA-MB-231 cells. WT or ERCC4-/- cells were cultured in DMEM containing varying concentrations of FEN1i (LNT-1) for 10 days. Viable cells were stained with Crystal-violet solution.
